# Safeguarding evolutionary history in a biodiversity hotspot: downscaling the EDGE metric to a regional application

**DOI:** 10.1101/2025.06.16.659859

**Authors:** Ignacio Ramos-Gutiérrez, Sebastian Pipins, Rafael Molina-Venegas, Mario Fernández-Mazuecos, Pedro Jiménez-Mejías, Juan Carlos Moreno-Saiz, Félix Forest

## Abstract

In the face of accelerating biodiversity loss, phylogenetically informed approaches offer critical insights for conservation planning from an evolutionary perspective. The EDGE (Evolutionarily Distinct and Globally Endangered) metric combines phylogenetic singularity with extinction risk to identify taxa that represent unique evolutionary history under threat. In this study, we focus on the global conservation relevance of angiosperms native to a regional biodiversity hotspot, the Iberian Peninsula. Our aim was to identify areas with a high concentration of EDGE species and to evaluate how effectively the current protected area network encompasses them.

Our findings reveal that threatened evolutionary history is primarily concentrated in mountainous and coastal regions. While several EDGE zones —areas containing unique and endangered evolutionary lineages— overlap with existing protected areas, particularly in mountains, others harbouring few but evolutionarily unique and highly threatened taxa remain largely unprotected.

This study highlights the value of applying global conservation metrics such as EDGE at regional scales. Our results provide a foundation for integrating evolutionary history into conservation prioritization in the Iberian Peninsula and offer a replicable framework for implementing the EDGE approach in other biodiversity-rich regions.

**Impact statement:** In the Iberian Peninsula, there are 22 EDGE zones that account for over 90% of angiosperm threatened evolutionary history.

## Introduction

Earth is currently undergoing a mass extinction event that, for the first time in the planet’s history, is being driven by a single species, humans (IPBES, 2019). More than 25% of all extant species are on the brink of extinction, a figure that increases to nearly 45% when considering flowering plants alone alone (Bachman et al., 2024; IUCN, 2023; Nic Lughadha et al., 2020). This critical situation urges conservation action aimed at minimizing the biodiversity loss caused by human activities, and hence the need for analytical tools that can optimise the trade-off between well-informed decisions and time and financial investment (Forest et al., 2015; Isaac and Pearse, 2018). A key component of conservation biology is prioritizing species and areas for conservation in what has been referred to as the “agony of choice” (Vane-Wright et al., 1991) or “Noah’s ark problem” (Weitzman, 1998). Often species prioritisations are based on subjective or difficult to quantify criteria (e.g. charismatic or flagship species, keystone species; (McGowan et al., 2020; Smith et al., 2012), while priority areas have typically been delimited based on species richness, rarity (e.g. narrowly distributed or endemic taxa), and threat status (Brooks et al., 2006; Darbyshire et al., 2017; IUCN, 2020; Myers et al., 2000; Sánchez de Dios et al., 2017). The availability of biodiversity data has vastly increased over the last decades (Wüest et al., 2020), providing new opportunities to assess priority species and areas in ways that better represent biodiversity’s value. For example, the rapid development of phylogenetic and genomic techniques has enabled the prioritisation of areas led by the amount of phylogenetic diversity they harbour. Phylogenetic diversity is a metric compiled by summing up the length of branches that connect a set of terminals on a phylogenetic tree (Faith, 1992). Similar approaches have been used such as prioritising species based on the fraction of unique evolutionary history they encompass (evolutionary distinctiveness; Redding and Mooers, 2006), a strategy that may in turn delimit priority areas based on the number of evolutionarily distinct species they harbour.

Isaac et al. (2007) developed a species prioritisation tool for conservation called the Evolutionarily Distinct and Globally Endangered (EDGE) metric. Rooted in the value-risk trade-off concept of economics (Isaac and Pearse, 2018; Weitzman, 1998), this metric combines information regarding the evolutionary history of species with their extinction risk, so that the highest priority for conservation is given to the most evolutionarily distinct (ED) and threatened (i.e. globally endangered, GE) species. Therefore, this metric can be used to effectively safeguard the most unique portions of the Tree of Life. The EDGE approach has been used to prioritize species across several major clades, including gymnosperms (Forest et al., 2018), chondrichthyans (Stein et al., 2018), and jawed vertebrates (Gumbs et al., 2024, 2018), and a priority list for angiosperms is currently in progress (Forest et al., 2025). Moreover, the framework has been incorporated into a conservation initiative that focuses on species deemed to be both threatened and evolutionarily distinct, the EDGE of Existence Programme (https://www.edgeofexistence.org/) and the EDGE index has been included as an indicator under Target 4 of the Global Biodiversity Framework (CBD, 2022). The EDGE approach involves, by definition, a global assessment of all known species in a clade of interest, which has lent itself towards global prioritisation studies (Pipins et al., 2024; Safi et al., 2013), though the framework is theoretically applicable to any spatial scale. Highlighting regional priorities of EDGE species can help to link national conservation efforts to global conservation goals, such as safeguarding the Tree of Life (Carta et al., 2019; Gumbs et al., 2023a). However, to the best of our knowledge, no regional EDGE analyses incorporating an entire flora have been conducted to date. Considering that conservation policy is typically implemented at country level, regional assessments of the geographical patterns of EDGE species may be instrumental for prioritizing effective and well-informed conservation actions that take into account the evolutionary dimension of biodiversity.

Here, we use the EDGE metric to conduct a spatial prioritisation at the regional level (hereafter, a regional EDGE assessment) for the angiosperm flora of the Iberian Peninsula in the western Mediterranean. This region is home to over 5,400 angiosperm species (over 20% of the Mediterranean angiosperm flora; Ramos-Gutiérrez et al., 2021; Rundel et al., 2016), one of the richest regions across the entire Mediterranean basin (Araújo et al., 2007). Moreover, it constitutes an exceptional centre of endemism (Buira et al., 2021; Cai et al., 2023; Médail and Quézel, 1999), thus representing an ideal setting for a first regional EDGE assessment. Specifically, we aim to identify areas of high conservation value due to the presence of a large number of EDGE species and provide a list of priority regions using a complementarity-based approach that sequentially identifies areas harbouring high-EDGE species across the region.

## Methods

### Study area

The Iberian Peninsula is located in the southwestern tip of Europe, occupying an area of approximately 590,000 km^2^. It encompasses the political territories of mainland Spain, mainland Portugal and Andorra, as well as the Balearic Islands, and a few minor offshore peninsular archipelagos (e.g. Berlengas, Cíes, Columbretes) that are geologically and floristically related to the Iberian Peninsula (Figure 1). This territory is crisscrossed by several mountain ranges that contribute to the floristic diversity and endemism of the region (Buira et al., 2017; Médail and Quézel, 1997). Most of the Iberian Peninsula has a Mediterranean climate, except for the northern and northwestern territories, where a temperate oceanic climate prevails.

**Figure 1:**
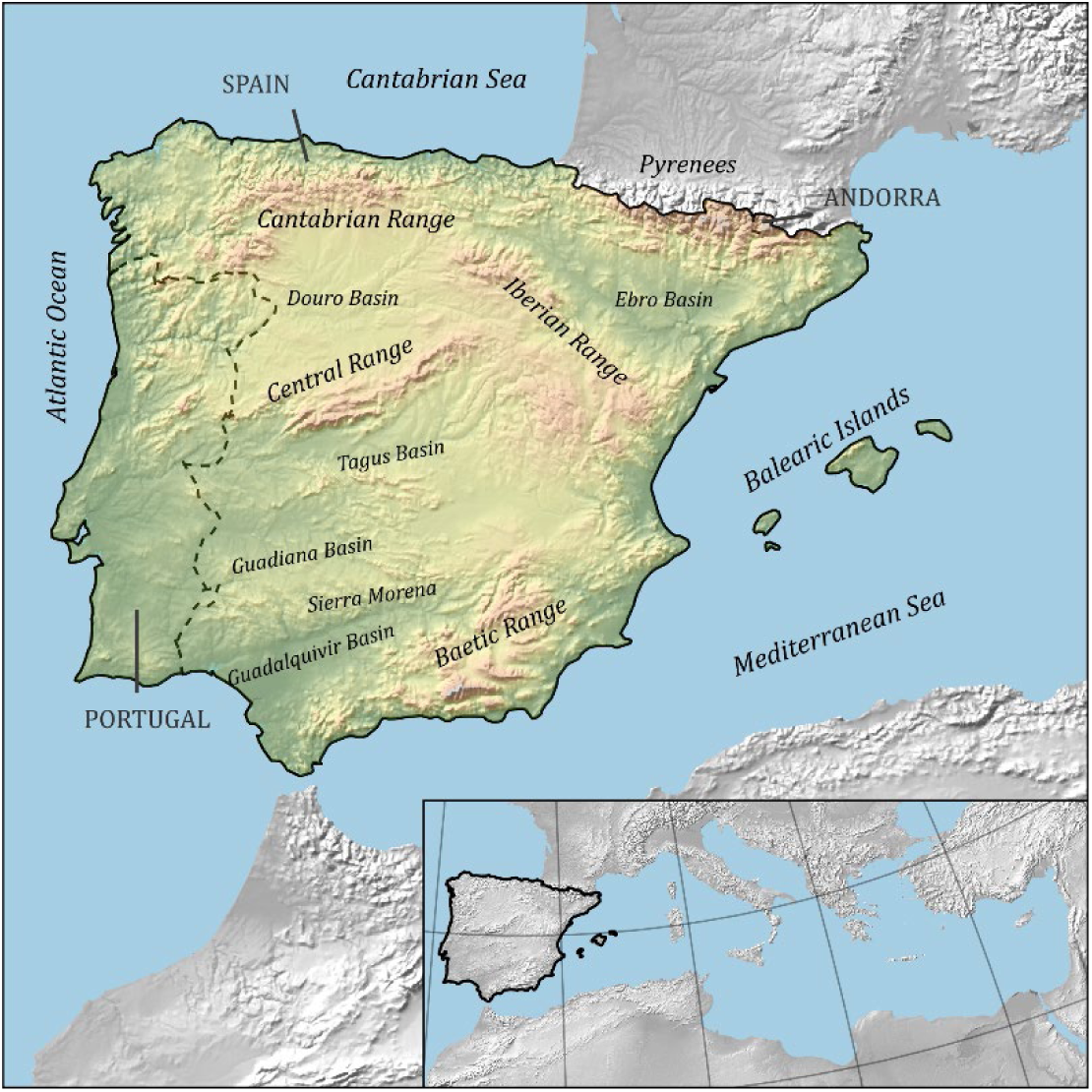
Topography of the Iberian Peninsula in the context of the Mediterranean basin (inset). The Iberian Peninsula (continuous contour) comprises mainland Spain and Portugal, Andorra, the Balearic Islands and several minor offshore islands (not shown). The political boundaries between countries (Andorra, Portugal and Spain) are indicated by dashed lines. Major mountain ranges and river basins are also depicted.

### Taxonomic and chorological data

We retrieved a global checklist of angiosperms from the World Checklist of Vascular Plants (WCVP version 11; Govaerts et al., 2021), comprising 335,497 species in total, and a checklist of angiosperm species native to the Iberian Peninsula from the AFLIBER database (Ramos-Gutiérrez et al., 2021). This resource documents the distribution of native vascular plant species in the Iberian Peninsula at 10×10 km resolution (totaling 6,303 UTM grid cells). To homogenize the taxonomic treatment between the sources, subspecific taxa were merged into their respective species, and names were homogenized to WCVP, resulting in a total of 5,411 Iberian angiosperm species.

### Phylogenetic data

We generated a set of phylogenies including all angiosperm species in the global checklist by expanding the seed plant phylogeny of Smith and Brown (2018)with the R package ‘V.Phylomaker’ (Jin and Qian, 2019). Missing species from the backbone tree (hereafter ‘phylogenetically unplaced taxa’, PUTs) were imputed within a randomly selected node descending from the crown node of its genus, or family in the case of missing genera. If the congenerics of a PUT formed a polyphyletic group in the backbone tree, the PUT was imputed to a randomly selected node within the largest cluster of the genus in the backbone tree. To account for phylogenetic uncertainty in subsequent analyses, this process was repeated 200 times to produce 200 different trees (see Forest et al., 2025 for details).

### Global extinction risk data

Information on the global extinction risk of angiosperms was retrieved from the IUCN Red List database (IUCN, 2021). For Spanish endemic taxa lacking a global assessment, we retrieved their status from the national Red List and the last update of the Spanish Red Data Book (Moreno Saiz, 2008; Moreno Saiz et al., 2019). For the rest of the species without formal IUCN Red List assessments (68.3%), we used the predictions of extinction risk from Forest et al. (2025), which designated species as either threatened or not threatened. The distribution of extinction risk categories for the Iberian angiosperm species can be seen in Table 1, while the spatial distribution of species included as threatened (i.e. Critically Endangered, Endangered, Vulnerable or predicted as threatened) in the Iberian Peninsula is shown in Figure S1.

**Table 1:** Distribution of global extinction risk categories across the 5,411 Iberian angiosperm species included in this analysis. The species assessed by the IUCN Red List are assigned to the categories EX (Extinct), EW (Extinct in the wild), CR (Critically Endangered), EN (Endangered), VU (Vulnerable), NT (Nearly Threatened), and LC (Least Concern), whilst the remaining species are treated as “threatened” or “non-threatened” species based on predictions of extinction risk. The global assessment column shows the number of taxa whose extinction risk was retrieved from Forest et al. (2025). The regional assessment figures stand for Spanish endemics whose category was upgraded based on Moreno Saiz et al. (2019). The last column, obtained by summing up the two previous, depicts the number of taxa for each category as used for the analyses.

| Extinction risk category | Number of species<br>(global assessment) | Number of species<br>(regional assessment) | Total number of species |
| --- | --- | --- | --- |
| EX | 1 | 1 | 2 |
| EW | 1 | 0 | 1 |
| CR | 30 | 62 | 92 |
| EN | 50 | 78 | 128 |
| VU | 46 | 162 | 208 |
| NT | 61 | 132 | 193 |
| LC | 691 | 398 | 1,089 |
| Other threatened | 285 | 0 | 285 |
| Other non-threatened | 3,413 | 0 | 3,413 |

### EDGE score calculation

EDGE scores are a quantification of how much threatened evolutionary history a taxon is responsible for. Thus, the more evolutionarily distinctive and/or threatened a species is, the larger the score it will receive. We calculated an EDGE score for each Iberian angiosperm species following the EDGE2 framework (Gumbs et al., 2023b). EDGE2 is an extension of the original EDGE framework that incorporates extinction risk data through a continuous distribution of extinction probabilities instead of discrete categories. Furthermore, the EDGE2 approach integrates information on the extinction risk of close relatives when calculating the ED2 score for a given species (Gumbs et al., 2023b), in which each internal branch of the tree is scored with a probability of extinction based on the combined extinction risks of all descendant branches. EDGE2 scores (hereafter, EDGE scores) were computed for each species across the 200 complete species-level phylogenetic trees following Forest et al. (2025), and the median value of these 200 EDGE scores per species were compiled for subsequent analyses. Species threatened with extinction (i.e. those assessed as EW, CR, EN or VU categories, or those predicted as threatened) or extinct (EX), and whose EDGE value was above the median EDGE score in at least 95% of the trees were identified as “EDGE species”, that is, priority species for conservation based on their evolutionary distinctiveness and global extinction risk.

### Spatial analyses

We calculated the threatened evolutionary history for each 10x10 km grid cell by summing up the EDGE scores of all the species present within that grid cell. Because summed threatened evolutionary history and species richness are expected to be intrinsically correlated, we also computed the mean EDGE score for species occurring in each grid cell. Subsequently, we identified priority areas for conservation in the Iberian Peninsula based on the complementarity of summed EDGE scores (Pipins et al., 2024; Vane-Wright et al., 1991). First, we identified the top-EDGE grid-cell across the study area, the one with the highest summed threatened evolutionary history, and all adjacent (i.e. sharing at least one side) and floristically similar grid-cells were clumped with the first-ranked cell. The rationale for this grid cell grouping is that selecting areas based on single grid cells may leave adjacent areas of high EDGE value unprotected due to incomplete sampling. Two grid cells were deemed floristically similar if their taxonomic dissimilarity, as measured by the turnover component of beta diversity (Baselga, 2010), fell below the 2nd percentile of the distribution of dissimilarity values between the focal cell and all the others. Then, all the species occurring in the resulting EDGE priority area were removed from the dataset, the next top-EDGE grid cell was identified, and adjacent and floristically similar cells were grouped as described above. This procedure continued iteratively until the complementary priority areas captured 90% of the total summed threatened evolutionary history of all Iberian angiosperm species. Pairwise beta-turnover between grid cells was computed with the beta.pair function of ‘betapart’ R package (Baselga et al., 2022). For each resulting EDGE zone we calculated the overlap with existing national parks, natural parks and natural reserves to get an estimation of its current protection status.

## Results

The Iberian Peninsula hosts 1,649 million years (My) of threatened evolutionary history. A total of 419 Iberian taxa were identified as EDGE species, representing 7.7% of all angiosperms in the region (see Table S1), and accounting for 501 My of threatened evolutionary history. These species belong to 50 families, with about half concentrated in only seven families, including Rosaceae (46), Brassicaceae (34), Fabaceae (30), Plantaginaceae (29), Caryophyllaceae (26), Poaceae (26) and Asteraceae (25) (see Figure S2). Most Iberian EDGE species are found in the mountainous regions of southern Spain (Figure 2b), yet the highest average scores (for EDGE species) are found along the inner, lower mountain ranges of Cazorla-Segura (Baetic Range) and Sierra Morena in southern Spain, with some points scattered throughout northwestern Spain and northern Portugal (Figure 2f). When considering the entire angiosperm flora (Figure 2e), highest prevalence of EDGE values are observed mainly in the northwestern corner of the Peninsula, indicating the presence of species not necessarily flagged as EDGE species but holding high EDGE values.

**Figure 2:**
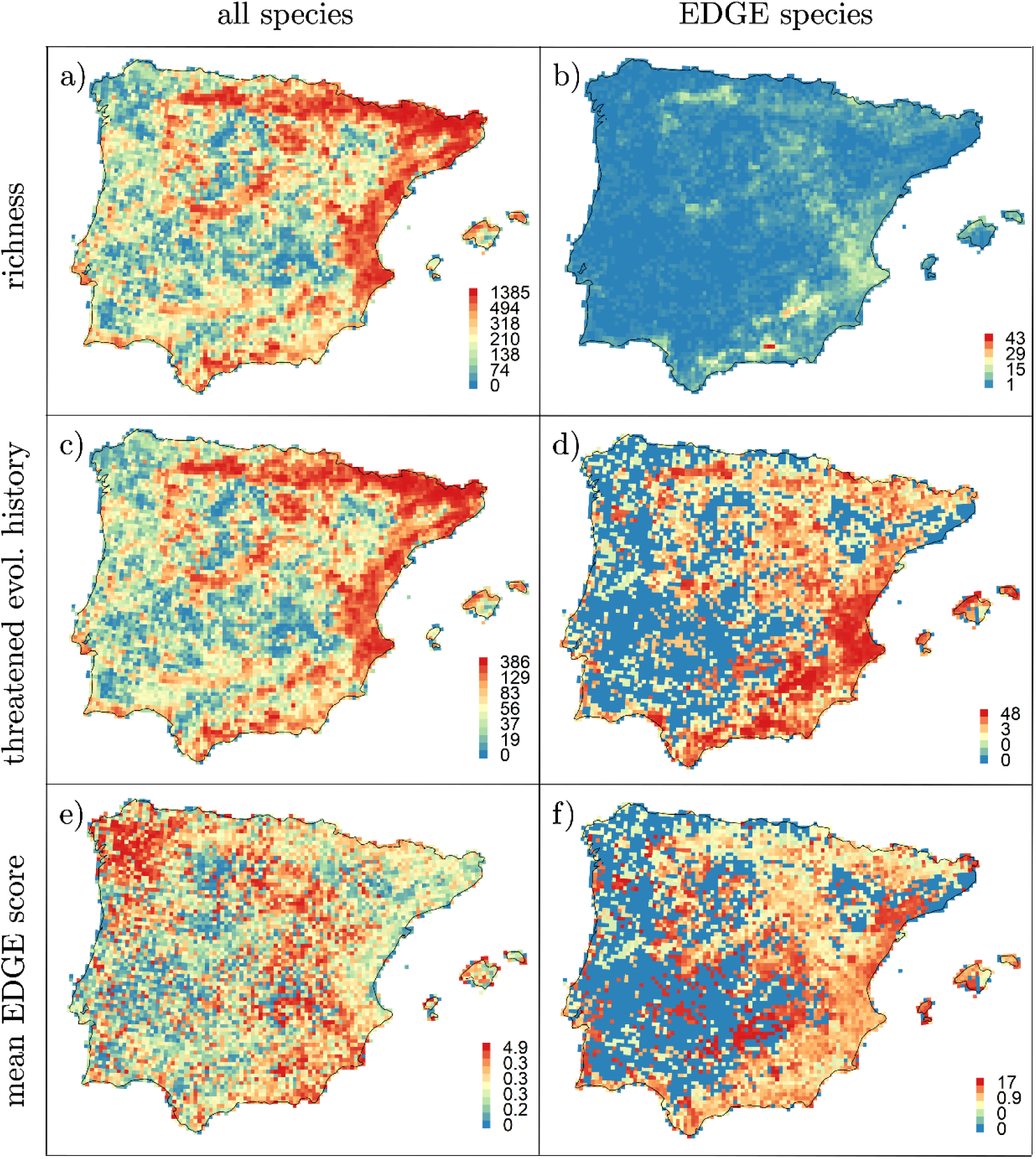
Distribution of EDGE metrics across the Iberian Peninsula for all angiosperm species (left column) and for the 419 taxa identified as EDGE species (right column). Species richness in each grid cell is shown in panels ‘a’ and ‘b’; panels ‘c’ and ‘d’ show the sum of EDGE values in each grid-cell (i.e. threatened evolutionary history, in millions of years), and panels ‘e’ and ‘f’ show mean EDGE score values.

We identified 22 priority areas, or ‘EDGE zones’ (Pipins et al., 2024; Safi et al., 2013; see Figure 3), which largely occurred along mountain ranges (e.g. EDGE zones 1, 2, 3, 5, 11, 14, 15, 16 and 18), coastal regions (zones 6, 12, 21 and 22) or islands (zones 4 and 19 in Majorca and Minorca islands, respectively), with few exceptions (e.g. zones 7, 8 and 17). The priority rank of each EDGE zone, the amount of threatened evolutionary history they captured, and the proportion of area that is already under national protection is shown in Table 2. Although most of the EDGE zones partially overlap with protected areas, only 17.8% of their total extent is currently under protection, and some zones (namely EDGE zones 8, 10 15 and 21) are not covered by national parks, natural parks or natural reserves. Nevertheless, they may be included to some extent in other regional or local protection figures, such as regional parks or European Natura2000 sites.

**Figure 3.**
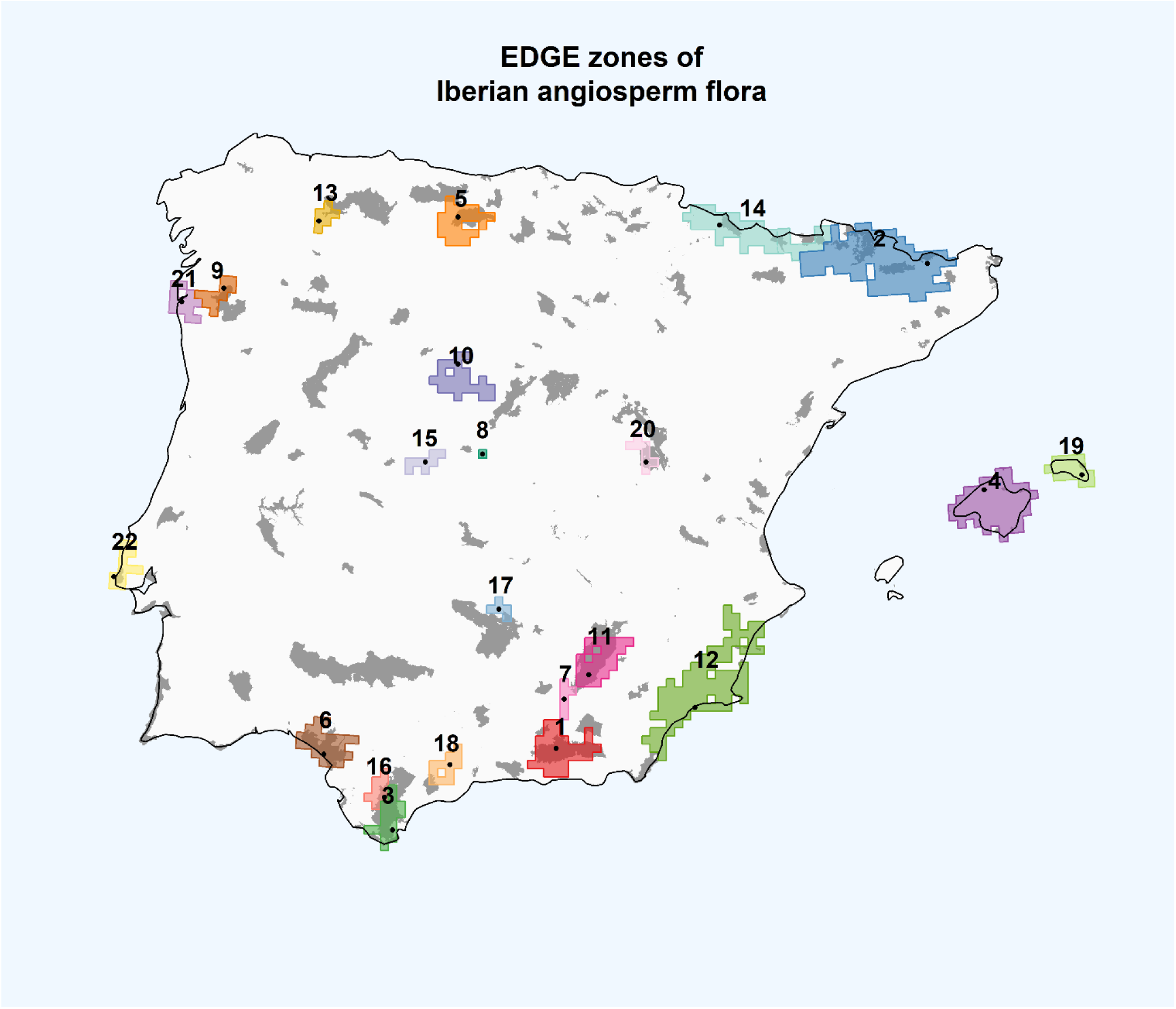
EDGE zones for the angiosperm flora of the Iberian Peninsula. The numbers labelling each EDGE zone match the numbers in Table 2 and Table S2. Black dots represent the selected grids in each EDGE zone. National parks, natural parks and natural reserves are shown in gray.

**Table 2.** List of EDGE zones and proposed names. The amount of threatened evolutionary history captured by each zone is also shown (in millions of years), as the cumulative proportion captured sequentially and the percentage of protected surface in each of them (under national park, natural park or natural reserve figures). The total percentage of national-management protection across all EDGE zones is 17.8%

| EDGE zone | Proposed EDGE zone name | Unique threatened<br>evolutionary history | Cumulative captured<br>evolutionary history | % Protected<br>area |
| --- | --- | --- | --- | --- |
| 1 | Sierra Nevada | 531 | 32.2 % | 49.5 % |
| 2 | Eastern Pyrenees | 376 | 55.0 % | 14.4 % |
| 3 | Campo de Gibraltar | 96 | 60.8 % | 69.19 % |
| 4 | Majorca | 68 | 65.0 % | 0.9 % |
| 5 | Central Cantabrian Range | 54 | 68.2 % | 28.8 % |
| 6 | Doñana | 39 | 70.6 % | 44.1 % |
| 7 | Hoya de Guadix | 31 | 72.5 % | 6.7 % |
| 8 | Cadalso de los Vidrios | 20 | 73.7 % | 0 % |
| 9 | Peneda-Gerês | 25 | 75.2 % | 22.5 % |
| 10 | Tierra de Pinares | 21 | 76.5 % | 0 % |
| 11 | Sierras de Cazorla y Segura | 33 | 78.5 % | 59.4 % |
| 12 | Levante Murciano-Almeriense | 51 | 81.6 % | 0.6 % |
| 13 | Sierra de Ancares | 16 | 82.6 % | 21.3 % |
| 14 | Central Pyrenees | 26 | 84.2 % | 12.3 % |
| 15 | Sierra de Gredos | 14 | 85.0 % | 0 % |
| 16 | Sierra del Aljibe | 13 | 85.8 % | 32.8 % |
| 17 | Valle de Alcudia | 10 | 86.4 % | 18.3 % |
| 18 | Sierra de las Nieves | 16 | 87.4 % | 7.0 % |
| 19 | Minorca | 17 | 88.4 % | 3.2 % |
| 20 | Serranía de Cuenca | 11 | 89.1 % | 44.7 % |
| 21 | Foz do Minho | 10 | 89.7 % | 0 % |
| 22 | Sintra-Cascais | 10 | 90.3 % | 12.0 % |

## Discussion

Halting biodiversity loss is a major commitment of the 2030 Agenda for Sustainable Development of the United Nations (goal 15; https://sdgs.un.org/goals), which demands effective conservation actions, particularly in biodiversity hotspots. The use of evolutionary based approaches can be useful for their consecution (De Meester et al., 2024), and thus to mitigate the biodiversity crisis we are currently witnessing. Here, we present the first regional EDGE assessment in one of these biodiversity hotspots, the Iberian Peninsula, marking a step toward identifying key areas in need of protection. Our evolutionary-based analysis can help regional Spanish and Portuguese policymakers in delimiting top-priority areas for the preservation of a huge amount of unique evolutionary history that is largely threatened. In the Iberian Peninsula, these areas of high conservation interest are distributed across the territory, often linked to coastal or montane regions, and most of them are poorly covered, or not covered at all, by officially protected areas. The limited overlap between high diversity areas and protected areas is consistent with previous studies conducted on vascular plants plants (Araújo et al., 2007; Castro Parga et al., 1996) and other biological groups in the Iberian Peninsula (Araújo, 1999; Lobo and Araújo, 2003; Sérgio et al., 2000) and elsewhere (Daru et al., 2019).

Species richness has been a major driver of the delimitation of important areas for biodiversity (Anderson, 2002; Plantlife, 2010; Sánchez de Dios et al., 2017), but new methodologies, including the EDGE approach, allow for a more refined integration across various components of biodiversity (Carta et al., 2019; Kling et al., 2018; Marchese, 2015). Phylogenetic diversity is one way of going beyond species richness measures (Faith, 1992). However, it can be combined with extinction risk information for more efficient biodiversity conservation policies (Isaac et al., 2007; Nee and May, 1997; but see Winter et al., 2013). In the study area, the richest areas for vascular plants are found in mountain ranges ranges (Castro Parga et al., 1996; Lobo et al., 2001; Ramos-Gutiérrez et al., 2021). Several explanations can be put forward to explain these patterns. These areas are heterogeneous in terms of micro-climatic and geomorphological features (Antonelli et al., 2018), hence offering a wide variety of habitats in relatively small areas that may allow the coexistence of functionally disparate plant lineages at the macroecological scale and acting as cradles for newly radiating species (Lobo et al., 2001; Rahbek et al., 2019). However, this environmental heterogeneity also drives a lack of a continuous suitable habitat for taxa at a local scale, which ultimately causes a high prevalence of specialized and restricted taxa, which are thus prone to be threatened (Hoorn et al., 2013). Ultimately, mountain regions in the area of study (both high and medium altitude ones) have been less impacted by human activities than the lowlands (Castro Parga et al., 1996), which may explain the persistence of disparate lineages and the concentration of high amounts of unique and threatened evolutionary history in these areas.

The compilation of the EDGE scores can highlight individual species on the basis of their evolutionary uniqueness and extinction threat, and can be spatially summarized to detect areas clustering EDGE species, and thus harboring high levels of threatened evolutionary history. The distribution of threatened evolutionary history (i.e. summed EDGE scores) in the Iberian Peninsula somewhat resembles patterns observed for species richness, with several mountain areas highlighted as EDGE species-rich regions (cf. Figure 2). This especially applies to the orogenic regions in southeastern Spain, namely the Baetic range and its extensions, including the Balearic Islands. The topographical characteristics of these ranges, with steeper reliefs and isolated alpine areas lead to the appearance of narrow-ranged and species, which are more likely to be threatened (Buira et al., 2021; Castro Parga et al., 1996; Davies et al., 2011; see Figure S1). The low mean EDGE score values observed throughout the Pyrenees and northeastern Mediterranean coast can be understood in the light of their low relative endemism values and thus relatively low number of globally endangered taxa thriving there (see Figure S1). This pattern opposite to the general richness and diversity trends observed in the rest of Iberian mountains, can be attributed to the great number of species shared with other neighboring temperate or alpine regions (such as the French Massif or the Alps).

Our set of priority areas consists of 22 unique regions whose preservation would secure more than 90% of the Iberian Peninsula threatened evolutionary history. These results align with previous studies regarding priority areas for plant conservation using different methodologies uninformed by phylogenies. Several of the EDGE zones match those considered by Castro Parga et al. (1996) and Sánchez de Dios et al. (2017). The first of these studies identified priority areas for conservation based on richness or weighted endemism, such as Sierra Nevada, central and eastern Pyrenees or Sierra de Algeciras, while the second prioritized the Cantabrian and Cazorla Ranges and the arid southeastern region based on several factors as threat statuses, rarity or taxonomic distances. All these areas were flagged as EDGE zones in our analysis. Some of them overlap with major protected natural areas, but only eight have more than 20% of their territory covered and only two have more than 50%. These zones with high levels of protection are mainly those in mountainous regions a bias which has already been described for other parts of the world (see Rouget et al., 2003 and references therein). A notable exception is Doñana (EDGE zone 6), whose protection is driven mainly by vertebrates (BOE, 1969), being a shrub-marshland territory which is home to the Iberian lynx and a hotspot of migratory waterbirds. However, our study has also identified priority areas which had not been highlighted previously, containing a small number of species that are highly distinct. Examples of these areas are Cadalso de los Vidrios (EDGE zone 8), home to *Gyrocaryum oppositifolium* (Boraginaceae), the species with the highest EDGE value of the Iberian Peninsula according to our analysis (see also Vargas et al., 2020), or Guadix (zone 7), where *Haplophyllum bastetanum* (Rutaceae) thrives. Additionally, Sierra de Gredos (zone 15) is recovered as an EDGE zone based on the presence of *Pseudomisopates rivas-martinezii* (Plantaginaceae). Although the latter zone appears as unprotected in our analysis, its inclusion within a regional-managed protected area (regional park) grants some level of protection. Overall, some of these areas (see also zones 13, 17, 20), have traditionally been overlooked in conservation programmes, as they are not necessarily species-rich regions. Given their limited number of focal taxa for conservation and the relatively small area they cover, these regions emerge as ideal scenarios for micro-reserve protection management (Laguna et al., 2001). Our results emphasize the importance of considering multiple dimensions of biodiversity, and evolutionary distinctiveness in particular, to prioritize conservation efforts.

Despite the growing need to implement evolutionary-based approaches in conservation planning (Faith et al., 2018; Owen et al., 2019), we acknowledge that there is room for substantial improvement in the use of EDGE as a key metric for conservation. Increasing information availability related to the conservation status of species (i.e. reducing the *Scottian* shortfall; Haelewaters et al., 2024) by updating and completing the IUCN Red List, including for example regional red list categories, would be instrumental in optimizing the results of EDGE analyses and improving their accuracy and reliability. A first logical step would be to prioritize the formal assessment of species that have a high EDGE score, but that have not yet been assigned a formal UICN Red List category (“EDGE Research List” *sensu* Gumbs et al., 2023b). Another major step forward would be increasing the representation of species in molecular phylogenetic analyses. The greater the number of phylogenetically placed taxa, the lower will be the phylogenetic uncertainty introduced by species missing molecular information (Ramos-Gutiérrez et al., 2023; Rangel et al., 2015). The production of DNA sequence data for unsampled species is therefore key to refine these analyses and other studies using metrics derived from large phylogenetic trees. Additionally, some light must be shed into the different possibilities of using the EDGE metric at a regional level, such as the use of national extinction risk assessments or a regionally pruned phylogenetic tree. However, the direct use of globally calculated metrics ensures that the results obtained in this study are consistent with the foundational sense of the EDGE metric, and respond to a global conservation responsibility (“*How much global evolutionary history can be protected in this region?*”), rather than a territorial conservation interest (“*How can the evolutionary history present in this region be better protected?*”). In spite of these limitations, we believe that the spatial patterns reported here are robust and unlikely to change radically, as they have been calculated using state-of-the-art methodologies and based on an up-to-date spatial dataset (AFLIBER) that stands out for its exceptional completeness and resolution, with few regional equivalents.

The priority set of regions outlined here may help guide managers to select important areas for the conservation of the flora they harbour. This task has already begun with the recent inclusion, among others, of the top-1 and top-9 ranked EDGE species in the Iberian Peninsula (i.e. *Gyrocaryum oppositifolium* and *Gadoria falukei*) in the list of Spanish protected taxa, after a requirement by the Spanish Plant Conservation Society (BOE, 2025; Moreno Saiz and Martínez García, 2023). This work represents a step forward in establishing synergies between phylogenetics, conservation biology, and policymaking at the regional level, and we hope that it will serve as a springboard for further application of the EDGE methodology in other regions of high biodiversity value.

## Acknowledgements

This work was possible thanks to the funding of the *Iberian Flora on the EDGE* project (TED2021-131234A-I0), led by M.F.M. and P.J.M. and the UAM-Santander Scholarships for young researcher mobility 2022, which covered living costs for a three month stay at RBG Kew. We would also like to acknowledge the James Hutton Institute for providing computational resources from the “UK’s Crop Diversity Bioinformatics HPC”.

---

**Figure S1:**
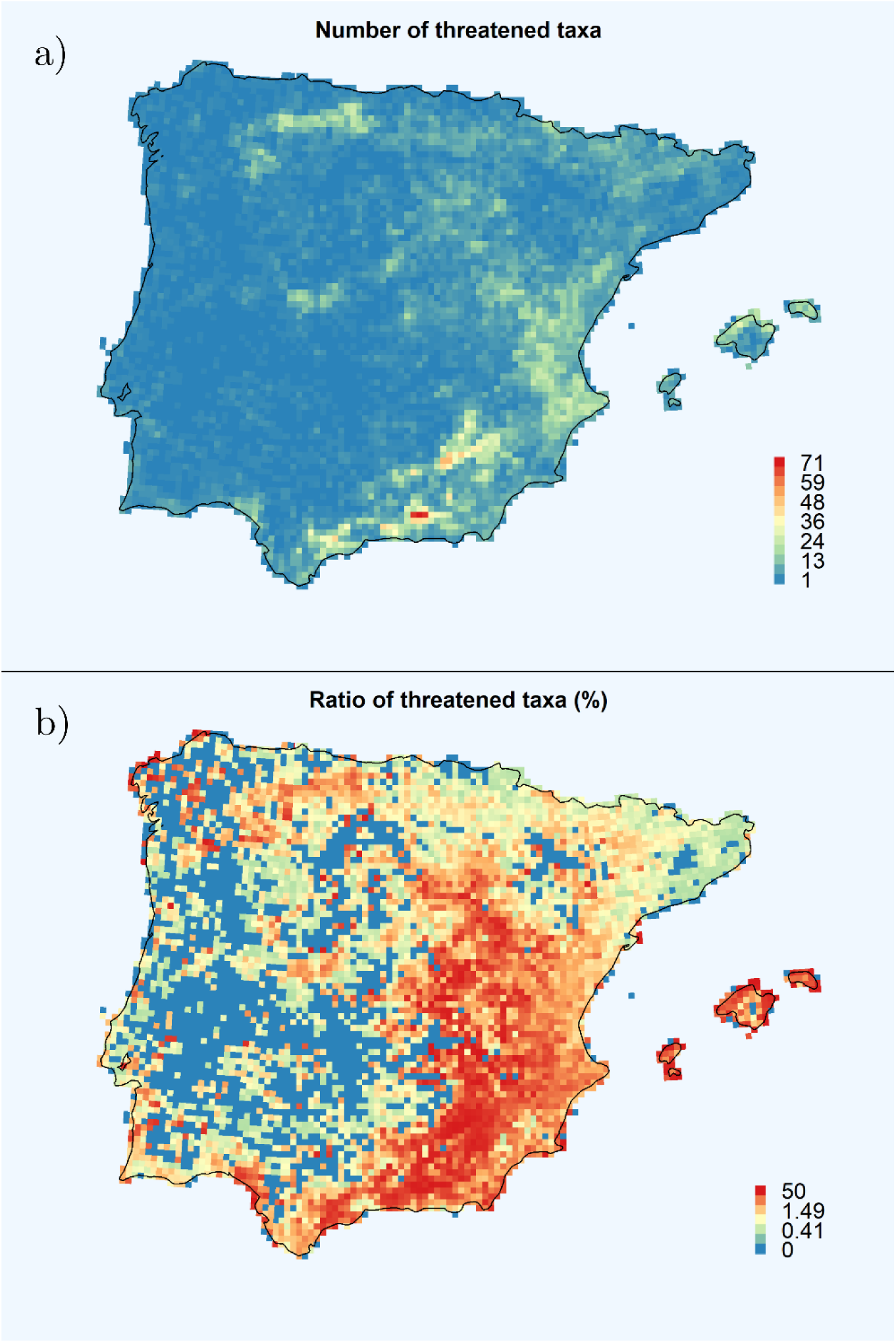
Spatial distribution of threatened species per grid cell in the Iberian Peninsula. Panel a) depicts the total number of threatened taxa (namely Critically Endangered, Endangered, Vulnerable and predicted as threatened) per 10x10km UTM grid cell; while panel b) shows the ratio of threatened taxa per site, calculated as the total number of threatened species divided by total number of species. Please note the colour palette stands for a non-parametric (equal-count) distribution for visualization purposes.

**Figure S2.**
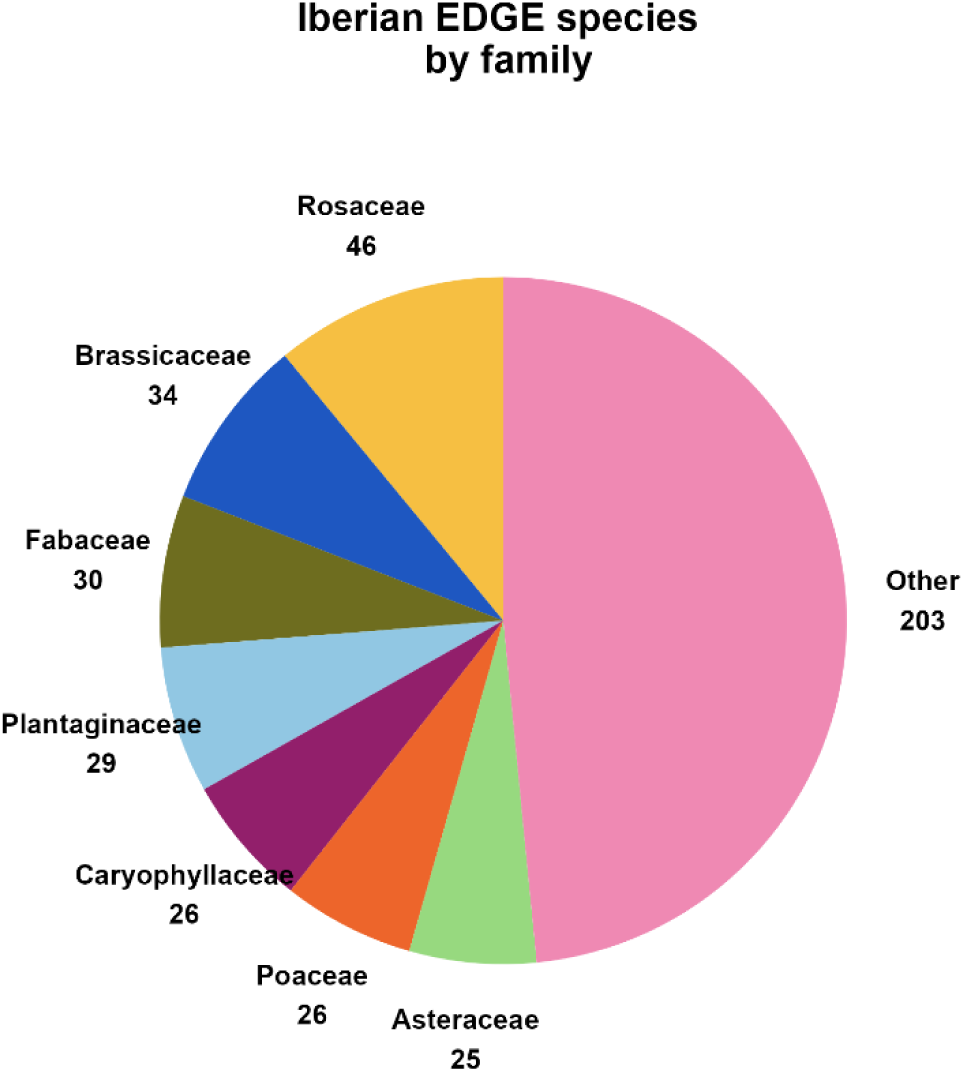
Proportion of Iberian EDGE species per family. Only the top 7 families (of a total number of 50) are shown, as they contain more than 20 EDGE species each, and over 50% of all the EDGE species.

**Table S1.**
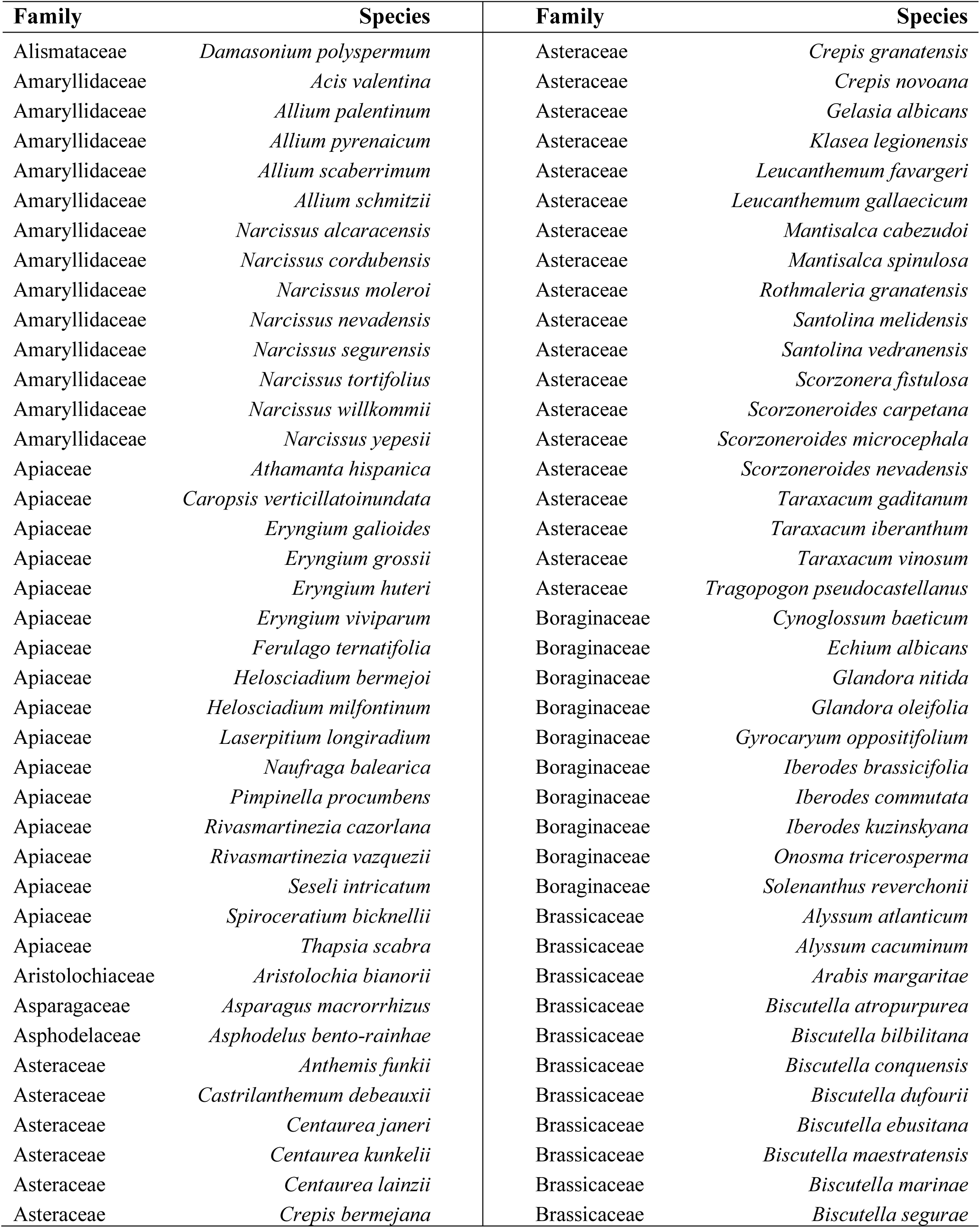
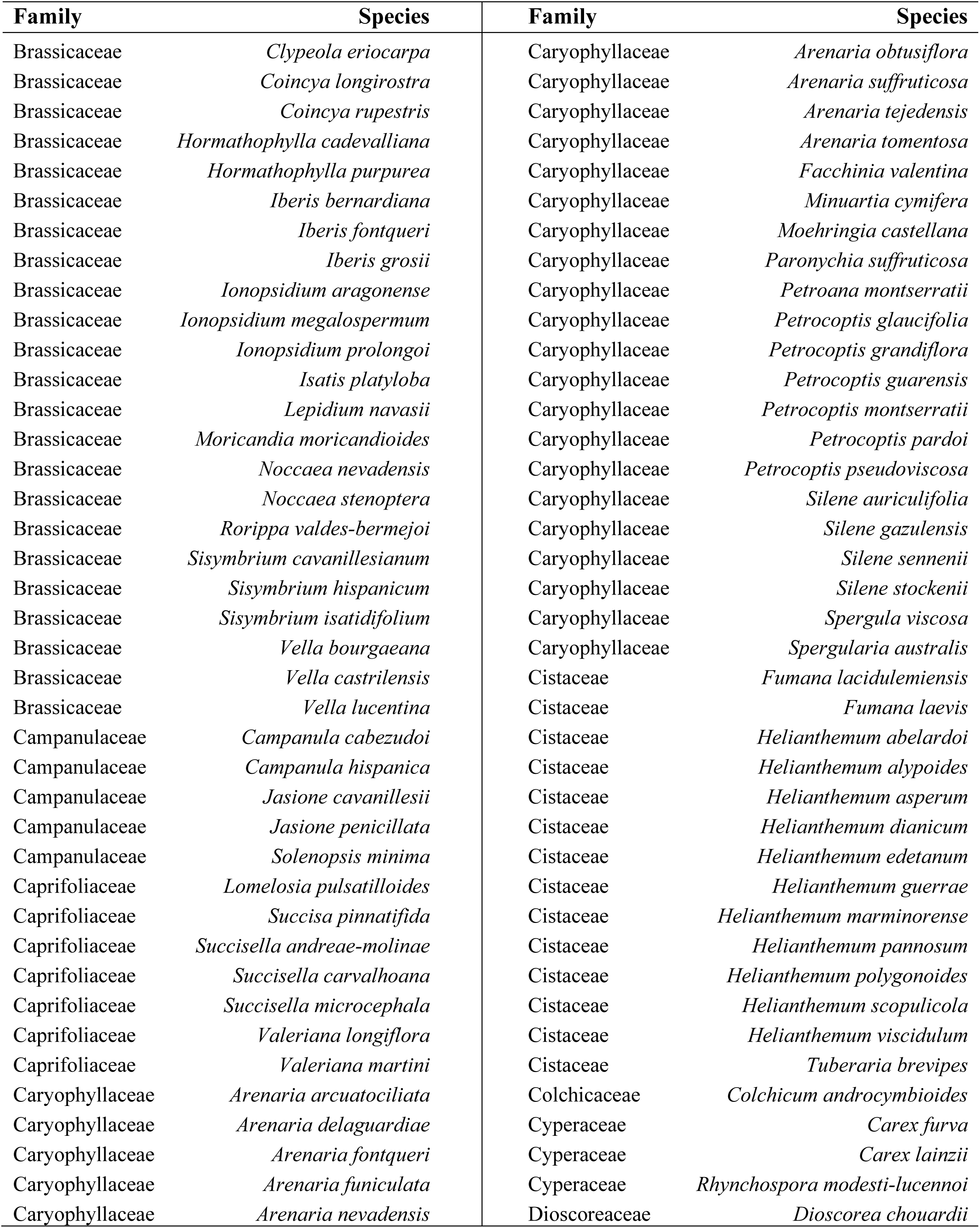
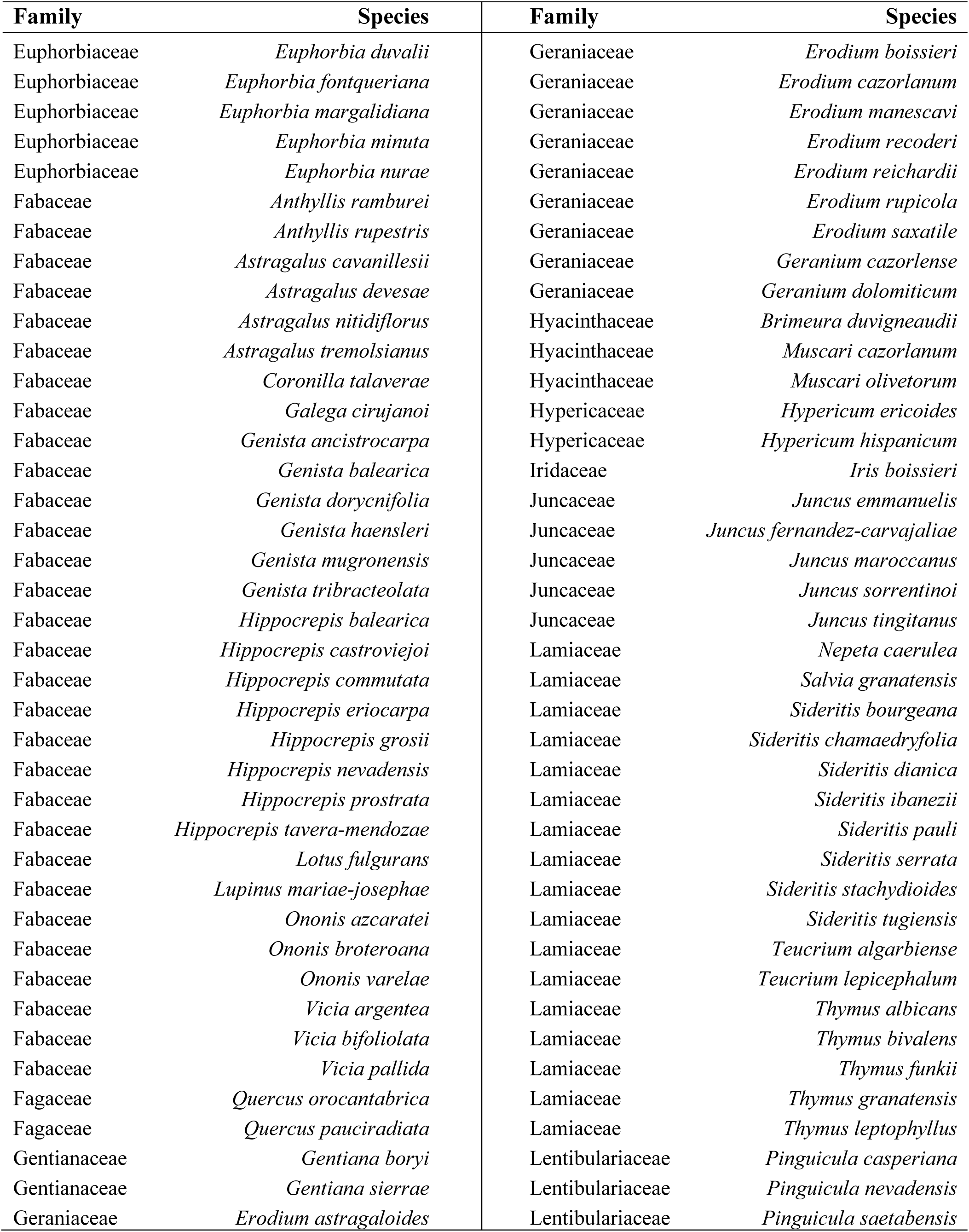
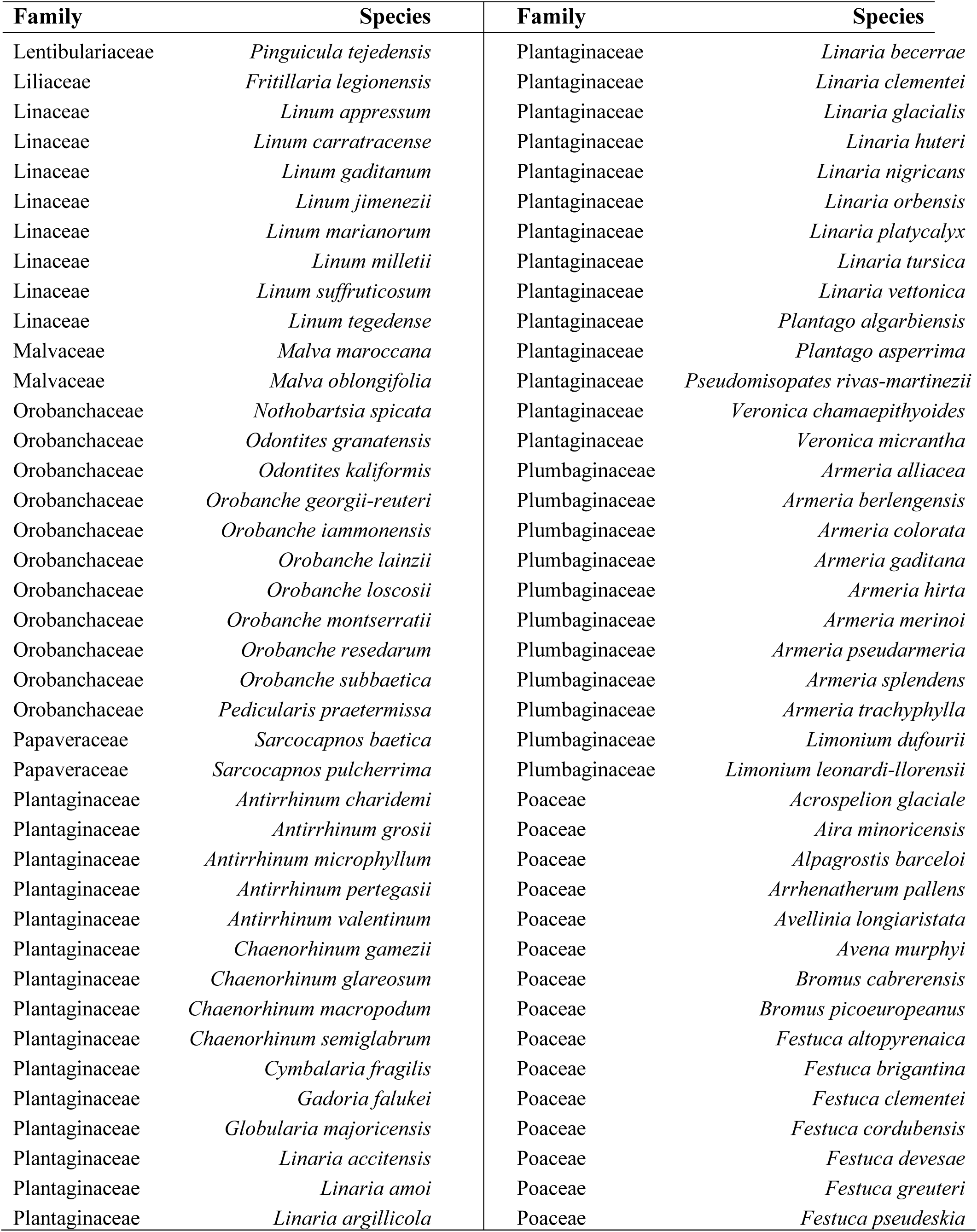
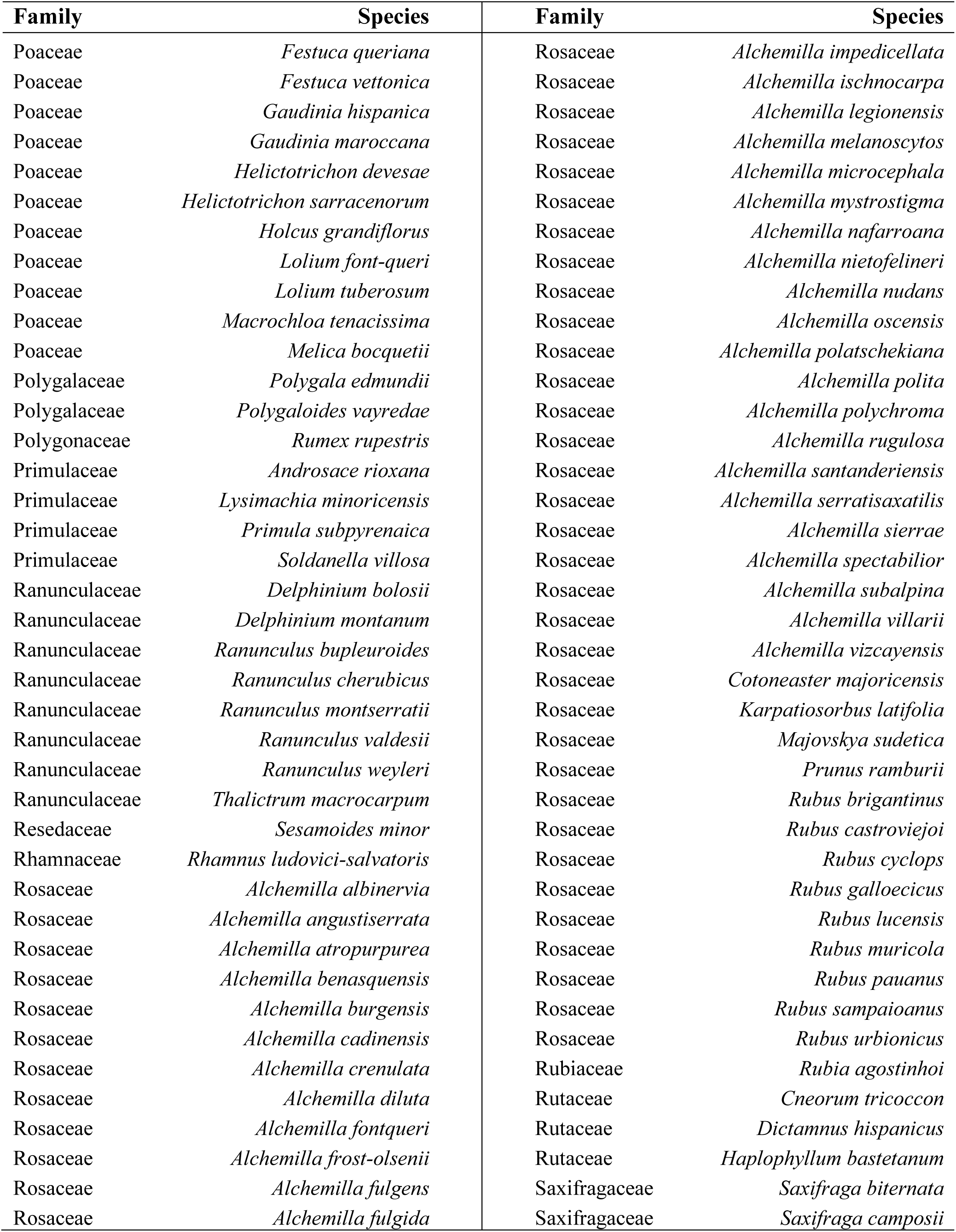
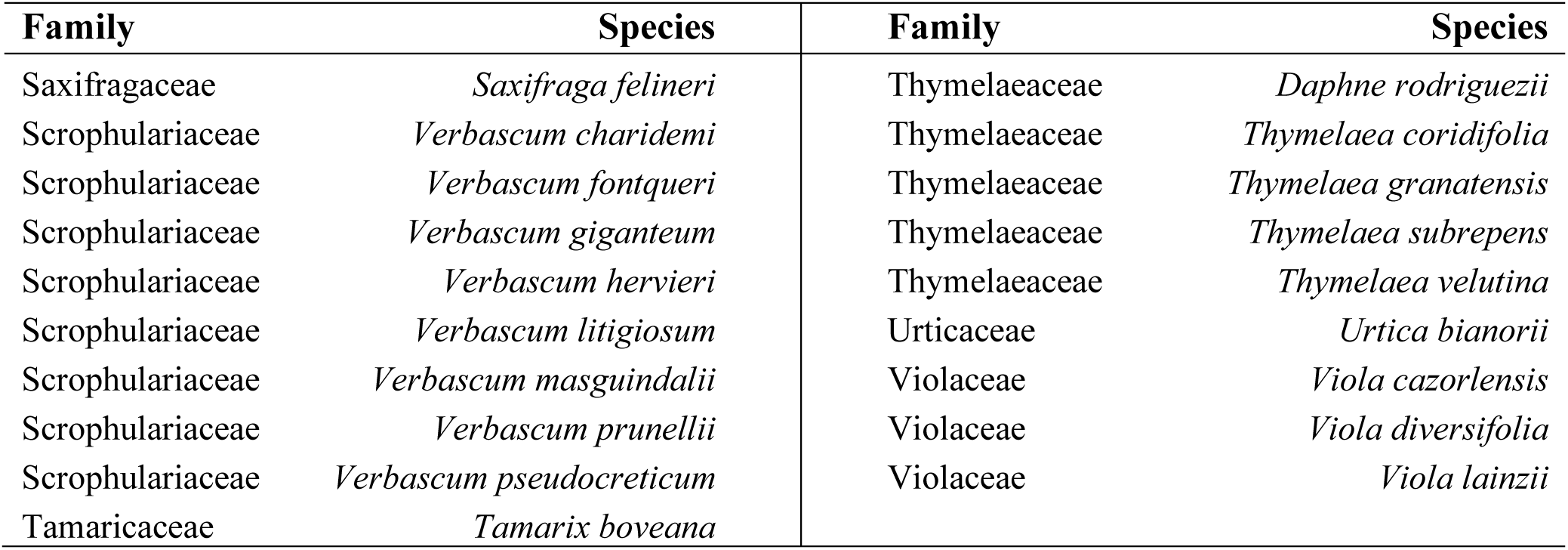
List of EDGE species and family to which they belong.

**Table S2.** Top-EDGE selected UTM 10x10 grid cells for each iterative step of the complementarity analysis and EDGE zone expanse (in square kilometers).

| EDGE zone | Selected grid cell | EDGE zone surface (km<sup>2</sup>) |
| --- | --- | --- |
| 1 | <u>30SVG70</u> | 3303 |
| 2 | <u>31TDG38</u> | 9904 |
| 3 | <u>30STF70</u> | 1699 |
| 4 | <u>31SDE80</u> | 5604 |
| 5 | <u>30TUN55</u> | 2400 |
| 6 | <u>29SQA29</u> | 2563 |
| 7 | <u>30SVG86</u> | 600 |
| 8 | <u>30TUK86</u> | 100 |
| 9 | <u>29TNG65</u> | 1100 |
| 10 | <u>30TUL57</u> | 2400 |
| 11 | <u>30SWG19</u> | 2602 |
| 12 | <u>30SXG45</u> | 8901 |
| 13 | <u>29TPH74</u> | 700 |
| 14 | <u>30TXN74</u> | 4857 |
| 15 | <u>30TUK15</u> | 800 |
| 16 | <u>30STF64</u> | 900 |
| 17 | <u>30SVH07</u> | 500 |
| 18 | <u>30SUF48</u> | 1200 |
| 19 | <u>31SFE01</u> | 1600 |
| 20 | <u>30TWK85</u> | 800 |
| 21 | <u>29TNG13</u> | 1301 |
| 22 | <u>29SMC59</u> | 1200 |

